# An introgressed galectin-like protein is a candidate driver of the human tropism in the intestinal parasite *Cryptosporidium*

**DOI:** 10.64898/2026.04.07.716958

**Authors:** Greta Bellinzona, Swapnil Tichkule, Aaron R. Jex, Cock van Oosterhout, Claudio Bandi, Davide Sassera, Michele Castelli, Simone M. Cacciò

## Abstract

*Cryptosporidium* spp. are protozoan parasites responsible for diarrheal diseases. In humans, cryptosporidiosis is predominantly caused by the human-specific *Cryptosporidium hominis* and by *Cryptosporidium parvum.* This second species has been classically reported as zoonotic, with a host preference for ruminants. However, the recently described subspecies *C. parvum anthroponosum* has been found to be restricted to humans.

Here, we generated novel whole genome sequences from West African samples of *C. p. anthroponosum*, and analyzed them together with all those already available, originating from East Africa, Europe, North America and Asia. Phylogenomics showed that all *C. p. anthroponosum* isolates are strongly clustered together, forming the sister clade of the zoonotic *C. parvum* representatives. The phylogenetic variations within *C. p. anthroponosum* did not present a clear geographic structure, consistent with *C. hominis*, primarily transmitted in humans.

To elucidate the evolution of host species adaptation in *C. p. anthroponosum*, we then investigated genetic exchanges with *C. hominis*, detecting an ancestral introgression present in all *C. p. anthroponosum* isolates. This introgression involved a single gene, encoding for an extracellular galectin-like protein, which we predicted with high confidence to form a protein complex with the human insulin-degrading enzyme, a key metabolic regulator. Considering the role of host insulin metabolism in the proliferation of parasites as well as its known intrinsic differences between humans and ruminants, this molecular interaction could represent a plausible mechanism for an important role of the galectin-like protein in host-parasite interactions and in the host specificity of *C. p. anthroponosum*.

## Introduction

*Cryptosporidium* is a genus of apicomplexan parasites causing diarrheal diseases in a broad range of vertebrate hosts. In humans, cryptosporidiosis is a leading cause of morbidity and mortality, particularly among young children living in low- and middle-income countries, where access to clean water and sanitation is limited ^1^. A recent estimate indicates cryptosporidosis as the cause of more than 48.000 deaths in children every year ^2^. However, the molecular mechanisms of interaction and pathogenicity are currently poorly understood, also hampering the development of effective treatment ^3^.

The genus *Cryptosporidium* currently comprises 44 species and over 120 genotypes, with marked differences in host specificity ^4^. Although many species are capable of infecting humans, most infections are due to *Cryptosporidium hominis* and *C. parvum*, classically considered anthroponotic and zoonotic, respectively. However, genetic data have revealed a more complex picture. Extensive molecular typing, in particular based on the highly polymorphic glycoprotein 60 (*gp60*) gene, disclosed the presence of many intra-species variants, or subtypes, which can also have different host preferences ^5^. More recently, this was corroborated by genomic analyses, which demonstrated the presence of both animal-adapted lineages in *C. hominis* ^6^ and human-adapted ones in *C. parvum* ^7^. In particular, the anthroponotic *C. parvum* subtype IIc was recognized as a separate subspecies, namely *C. parvum anthroponosum*, distinct from the zoonotic *C. p. parvum*. *C. p. anthroponosum* was suggested to have undergone adaptive divergence from zoonotic ancestors ^7^. Nevertheless, the small sample size and limited geographic representation of the available whole genome sequencing (WGS) data at the time, comprising only three *C. p. anthroponosum* samples from the United Kingdom, hindered a detailed reconstruction of the genomic and evolutionary processes underlying adaptation to the human host.

Here, we generated novel WGS data of *C. p. anthroponosum* from Western African samples, and leveraged all the current publicly available sequences of this subspecies, which allowed us to perform an in-depth phylogenomics and comparative genomics study with extended geographical coverage, spanning four continents. Thus, we identified an introgressed gene from *C. hominis* encoding for a galectin homolog, which was predicted to bind to a human insulin-degrading enzyme, thus providing potential evolutionary and functional underpinnings for the human tropism of *C. p. anthroponosum*.

## Materials and Methods

### Oocyst purification, DNA extraction and sequencing

Fecal samples had been collected during a study on transmission pathways of cryptosporidiosis among African children ^8^. By *gp60* sequencing, three samples collected in Ghana (AFR10297, AFR20297 and AFR16339) were shown to belong to the anthroponotic lineage IIc (subtype IIcA5G3a), thus were selected for the present study. The procedures for oocyst purification and DNA extraction were detailed in Corsi et al. (2023)^9^. In short, oocysts were purified from fecal specimens by immunomagnetic separation, treated with bleach, and used for genomic DNA extraction. Genomic DNA was then subjected to whole-genome amplification (WGA) using the REPLI-g midi kit (Qiagen), according to the manufacturers’ instructions.

For high-throughput sequencing experiments, ∼1 μg of purified WGA product per sample was used to generate Illumina Nextera XT 2×150 bp paired-end libraries, which were sequenced on an Illumina NovaSeq 6000 SP platform. Library preparation and sequencing were performed by a commercial service (Biodiversa, Italy).

### WGS dataset

Genomic data from additional *Cryptosporidium* samples were selected from NCBI for genomic analyses (Detailed sample information is provided in Supplementary Table S1). These comprised all available (n=11) *C. p. anthroponosum* samples (IIc subtype) ^7,10,11^, 13 representative *C. p. parvum* (IIa and IId subtypes, plus IIc-j)^7,9,10,12,13^ and 26 representative *C. hominis* samples ^14–17^ from diverse geographic regions, and a *C. meleagridis* sample ^14^, included as an outgroup. Raw sequencing reads were downloaded from the NCBI Sequence Read Archive (SRA) for all these samples and used for the following analyses.

### Raw data quality check and SNP calling

All raw sequencing reads were assessed for quality and processed with fastp v. 0.23.2 ^18^ to remove adapters and low-quality bases, using default settings with automatic adapter detection enabled. Trimmed reads were subsequently aligned to the *C. parvum* IOWA-ATCC reference genome ^19^ using minimap2 v.2.22 ^20^ with default settings.

Variant calling was performed following the standard GATK v.4.6.1.0 pipeline ^21^. Accordingly, BAM files were sorted by chromosome using Picard SortSam, and duplicates were identified and marked with Picard MarkDuplicates to improve the accuracy of variant detection. Average coverage for each sample was checked using samtools depth ^22^ and verified to be above 20X. Haplotype calling was performed using GATK HaplotypeCaller ^21,23^ in GVCF mode. SNPs were removed in case of quality depth <2.0, Fisher strand >60.0, mapping quality <30.0, mapping quality rank-sum test <−12.5, read position rank-sum test <−8.0, or strand odds ratio >3.0. Individual GVCF files for each sample were subsequently imported into a GenomicsDB workspace using GATK GenomicsDBImport. Finally, all single VCFs were merged using GATK GenotypeGVCFs to produce a merged VCF file. SNPs were further filtered using bcftools v.1.19 ^24^ based on the following criteria: biallelic SNPs only, quality score >30, allele depth >20, minor allele frequency >0.1, and missing data ratio <0.5.

To assess the multiplicity of infection (MOI) within samples, we applied MOIMIX (https://github.com/bahlolab/moimix) to calculate the Fws statistic, a fixation index used to evaluate within-host genetic differentiation. Following established guidelines ^25^, all samples were treated as clonal and kept for the following analysis, having Fws ≥ 0.95.

### Phylogenomics

A concatenated set of genomic SNPs was created by converting the VCF into a FASTA file with vcf2phylip v.2.0 ^26^ using the option “-f”. Then, a maximum-likelihood phylogenetic tree was inferred using IQ-TREE2 v.2.0.3^27^ with 100 bootstrap replicates (-b). The best-fit substitution model, namely TMV+F+ASC+R2, was automatically selected using ModelFinder^28^ with ascertainment bias correction.

### Recombination analyses

To explore potential recombination patterns, phylogenetic networks were first reconstructed using SplitsTree App ^29^, which allows visualization of conflicting phylogenetic signals and reticulate evolution.

Afterwards, the ABBA-BABA test ^30^ was performed with a dedicated set of scripts (https://github.com/simonhmartin/genomics_general), for which taxa were defined as follows: P1 is *C. p. parvum*, P2 is *C. p. anthroponosum*, P3 is *C. hominis*, and the outgroup is *C. meleagridis*. Accordingly, the fD statistic was calculated using default parameters in sliding windows of 5 kb with a 2.5 kb overlap, and considering only those windows containing at least 100 SNPs.

Only windows exceeding the 95^th^ percentile (i.e., top 5%) of the fD distribution (fD > 0.08) were further considered. For each of those windows, a maximum-likelihood phylogeny was reconstructed using IQ-TREE2 as described above.

Based on such analyses (see Results and Discussion), a putative introgressed genomic region was selected, to be further inspected and visualized using HybridCheck v.1.0.1 ^31^. The nucleotide and amino acid consensus sequence of the gene encoded in the candidate introgressed region was reconstructed from variant call format (VCF) data with respect to the reference sequence in the Iowa-II ATCC using GATK FastaAlternateReferenceMaker ^21^.

Furthermore, the gene annotated in this region was analysed with Hyphy v.2.5.8 using the Genetic Algorithm Recombination Detection (GARD)^32^, in order to further refine the boundaries of the introgression.

### Functional and evolutionary predictions on the candidate introgressed gene

Selective pressures on the single candidate introgressed gene sequence (see Results and Discussion) were evaluated using the HyPhy Branch-Site Unrestricted Statistical Test for Episodic Diversification (BUSTED)^33^, which tests for evidence of gene-wide episodic positive selection.

The consensus sequence of the protein encoded in the candidate introgressed region of *C. p. anthroponosum* UKP13, previously reconstructed from VCF data relative to the sequence (QOY42690.1) in the reference genome, was used for subsequent analyses. The Deeploc2 server (accessed in October 2025)^34^ was then used to predict the subcellular localization of this protein.

### Protein structure and interaction predictions

The three-dimensional structure of the consensus sequence of the introgressed protein in *C. p. anthroponosum* UKP13 was predicted using the AlphaFold3 server ^35^, then compared against known protein structures using the Foldseek webserver (accessed in October 2025) ^36^ to identify potential homologs.

To identify potential interactions with the human proteome, 15,187 human proteins expressed in the intestine were retrieved from The Human Protein Atlas^37^ (accessed December 2025), and the respective canonical protein sequences were downloaded from UniProt^38^. The potential interaction of the *C. p. anthroponosum* protein with each of those human proteins was tested using AlphaPulldown v2.0.1^39^, using MMseqs2^40^ to generate multiple sequence alignments. Initially, following the procedure described by Bellinzona et al. (2024) ^41^, the “--model_names=model_2_multimer_v3” option was used to generate a single protein complex model per each tested partner. Then, protein complexes with a pDockQ score greater than 0.5, considered a sufficiently stringent threshold ^42^, were re-assessed with AlphaPulldown with options “--num_predictions_per_model=5” and “--num_recycles 3”, thus generating 25 models for each complex, to increase specificity. Only complexes with a final pDockQ score above 0.5 were retained.

Visualization of protein and protein-complex structures were performed using ChimeraX v.1.11.1 ^43^.

## Results and Discussion

### *Cryptosporidium parvum anthroponosum* is a deeply divergent, human-associated monophylum

*C. p. anthroponosum* was recently characterized as a human-specific parasite ^7^, yet the genetic and molecular bases of this host tropism remain unknown. Here, we aimed to investigate the genomic evolution of *C. p. anthroponosum*, with a focus on identifying distinctive genetic features with respect to the zoonotic *C. p. parvum* and understanding their functional links to adaptation to humans. To this end, we newly sequenced the genome of three *C. p. anthroponosum* from Ghana, the first ones from Western Africa, and analyzed them together with all publicly available WGS (n=11) of this subspecies from Europe, Asia, North America and Africa, as well as a set of geographically and genetically diverse *C. p. parvum* (n=13) and *C. hominis* (n=26). Mapping the WGS reads to the *C. parvum* IOWA-ATCC reference genome led to the identification of 181,960 high-quality biallelic SNPs in this dataset.

The phylogeny reconstructed from this large set of biallelic SNPs revealed a fully supported deep split between the *C. parvum* and *C. hominis* lineages (Figure 1). In addition, within *C. parvum*, *C. p. anthroponosum* (subtype IIc) and *C. p. parvum* (subtypes IIa and IId, plus IIc-j) formed two distinct and fully supported sister clades. Therefore, the expanded taxon and geographic sampling strongly support that the two subspecies represent distinct evolutionary lineages within *C. parvum*, rather than being simply subtype variations. Interestingly, based on branch lengths, the genetic divergence between these two subspecies appears markedly larger than that between the two main *C. hominis* subclades, which were also recently proposed as subspecies ^44^.

**Figure 1.**
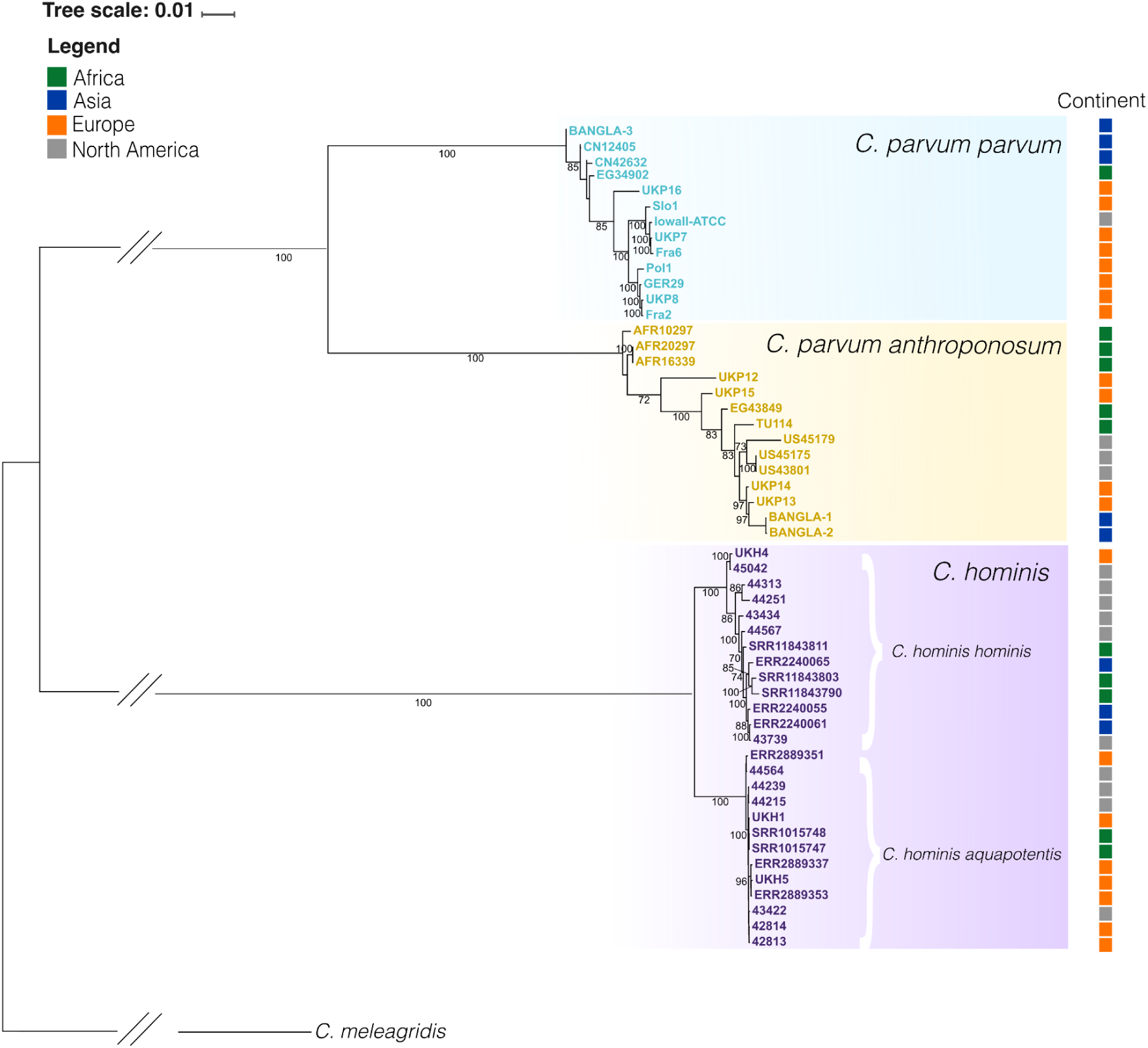
Maximum-likelihood phylogeny of *Cryptosporidium* samples based on over 181,000 high-quality biallelic SNPs. *C. p. parvum* samples are shown in blue, *C. p. anthroponosum* in yellow, and *C. hominis* in purple. *C. p. anthroponosum* forms a monophyletic highly-supported sister clade to zoonotic *C. p. parvum*, while *C. hominis* clusters separately. The colored squares on the right hand side indicate the geographical origin of each sample (legend on the top left). Only bootstrap values ≥70 are displayed. Tree scale stands for estimated proportional divergence. Double slashes indicate branches that were shortened for improved readability.

The monophyly of *C. p. anthroponosum*, together with its consistent detection exclusively in humans ^7^, is strongly indicative of an evolutionary conserved human tropism for this lineage. This is also supported by the lack of a clear geographic subdivision within this subspecies, even across continents, which is reminiscent of the structure observed in *C. hominis* ^44^ and contrasts with the evident geographic structure within *C. p. parvum*, with all European and North American isolates clustering together, as previously described ^12^.

### A localized introgression between *Cryptosporidium hominis* and *Cryptosporidium parvum anthroponosum*

Then, we sought to investigate the genomic and genetic processes at the basis of the host specialization of *C. p. anthroponosum*. To this end, considering that genetic exchanges with *C. hominis* were previously implied in the emergence of human tropism in *C. p. anthroponosum* ^7^, we focused on detecting such kinds of introgression events. A genome-wide split network analysis indicated the presence of appreciable, though not extensive, gene flow between *C. hominis* and *C. p. anthroponosum* (Supplementary Figure S1). To localize introgressions, we performed the ABBA–BABA test. Since fD>0 values were indicative of events involving *C. hominis* and *C. p. anthroponosum* (Figure 2A), we selected the genomic windows with the top 5% fD values (fD > 0.08; red dashed line in Figure 2B) as the most promising candidates for introgression, and evaluated each of them by inferring the respective phylogeny. Remarkably, the two overlapping windows (chromosome 3: position 840,001 to 847,500) that presented by far the highest fD (both >0.35, as compared to <0.19 for all others; Figure 2B) also displayed a tree topology consistent with an introgression from *C. hominis* to *C. p. anthroponosum*, namely a close proximity of the latter two with high support (Supplementary Figure S2). These results strongly indicate the occurrence of an ancestral introgression from *C. hominis* underlying the entire *C. p. anthroponosum* clade, rather than recent events restricted to individual isolates.

**Figure 2.**
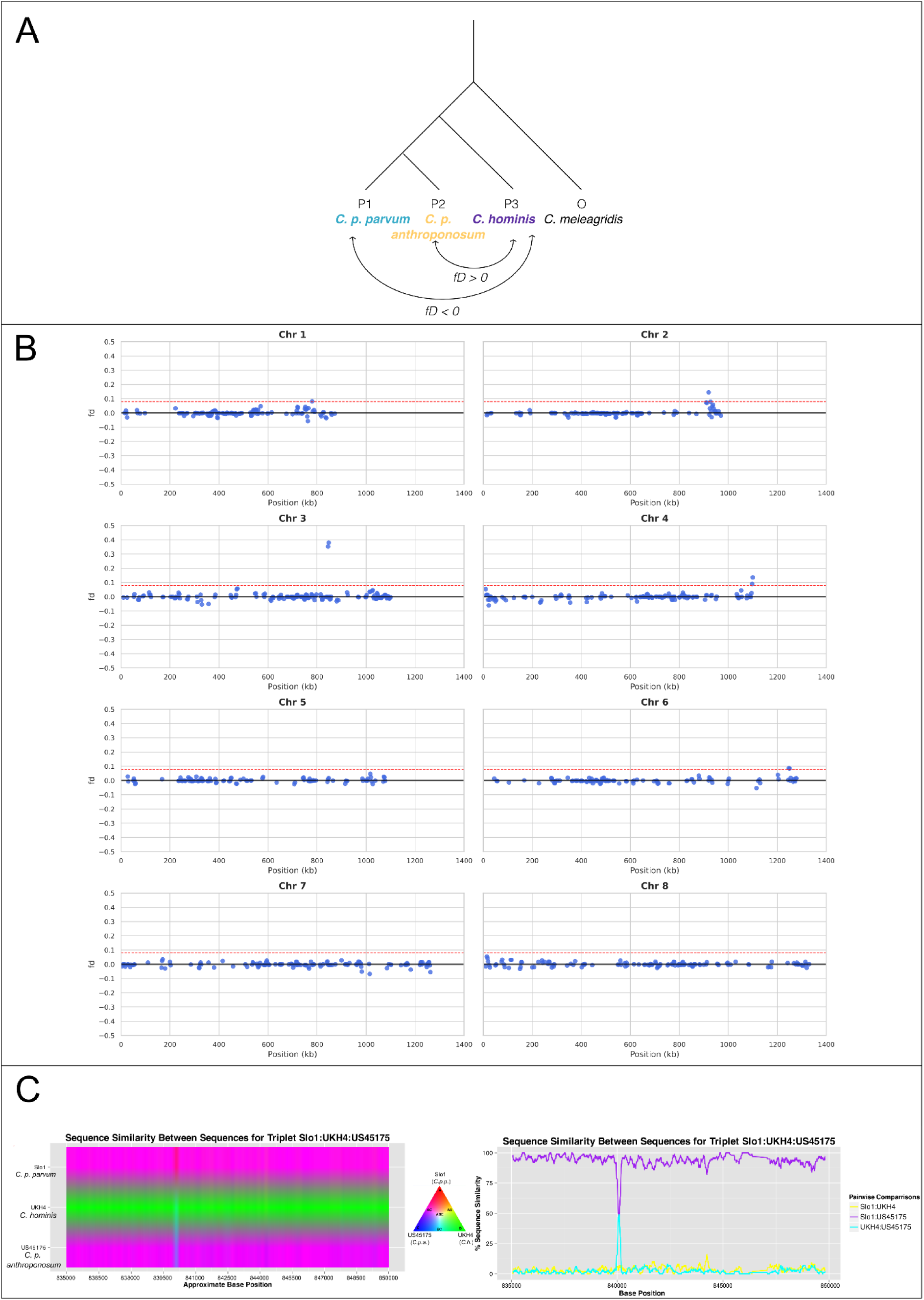
Analyses of introgression between *C. hominis* and subspecies *C. parvum*. A) Scheme of the lineages used for the genome-wide ABBA–BABA analysis, testing for localized introgressions between *C. hominis* (P3) and each *C. parvum* subspecies, namely *C. p. parvum* (P1) and *C. p. anthroponosum* (P2), with *C. meleagridis* as outgroup. Resulting fD values close to 0 indicate no detectable introgression, positive values indicate potential introgression between P2 and P3, while negative values indicate those between P1 and P3. B) Per chromosome fD statistic computed on 5,000 bp windows with 2,500 bp overlap. The red dashed line (fD = 0.08) indicates the threshold used to identify candidate introgression events. C) HybridCheck analysis of the chromosome 3 candidate region using three representative isolates: UKH4 (*C. hominis*), US45175 (*C. p. anthroponosum*), and Slo1 (*C. p. parvum*). The plot of the left represents the similarity between the three isolates, each in a row. Along the length of the region, colors represent whether the local SNPs are specific to the given isolate (primary color, respective vertex in the triangle in the middle) or to which degree they are shared with either of the other two isolates (color shades along the triangle sides). The plot on the right presents the pairwise similarity among isolates.

### The single gene encoded in the introgressed region is an extracellular structural homolog of galectins

We then conducted further analyses to confirm the introgression and define the boundaries of the genomic region involved. Using HybridCheck ^31^, we identified a peak of similarity between *C. hominis* and *C. p. anthroponosum* around position 840,078 of chromosome 3 (Figure 2C), while the GARD ^32^ algorithm suggested potential breakpoints at positions 841,475 and 843,100. According to the *C. p. parvum* IOWA reference genome annotation, a single coding sequence spans over the thereby refined region of introgression (accession: QOY42690.1, position 839,521 to 844,044). This gene did not show statistically significant evidence of episodic positive selection in the analysed *Cryptosporidium* dataset (BUSTED, p = 0.4425). However, this does not argue against an adaptive role of the introgressed haplotype. Namely, if host tropism changed through acquisition of an already divergent allele from *C. hominis*, the crucial evolutionary event may have been the introgression and retention of that intact haplotype, rather than a subsequent burst of amino-acid substitutions within *C. p. anthroponosum*. Under this scenario, little or no elevation in dN/dS within *C. p. anthroponosum* would be unsurprising. In the reference genome, the single protein encoded in the introgressed region (1059 amino acids) is labeled as a hypothetical uncharacterized protein. Database searches on NCBI nr yielded only a limited number of homolog sequences, all corresponding to equally uncharacterized proteins from *Cryptosporidium* spp., including *C. hominis*, *C. meleagridis*, *C. mortiferum*, *C. ubiquitum*, *C. canis*, and *C. felis*. To investigate the function of this protein, we predicted its three-dimensional structure, detecting the presence of two domains (residues 410–720 and 970–1303, highlighted in orange in Figure 3A), with strong structural similarity to a *Mus musculus* galectin (UniProt ID: Q9JL15; probability = 1, E-value < 0.01). We will refer to them as galectin-like domains. Galectins are extracellular proteins involved in glycan recognition, widely implicated in host–pathogen interactions, where they mediate processes such as cell–cell adhesion, host cell recognition, immune modulation, and surface attachment ^45,46^. Consistent with the homology with galectins, the *C. p. anthroponosum* galectin-like protein was also predicted to be extracellular. A large part of the amino acid sequence is identical across all *C. p. anthroponosum* samples as well as among *C. hominis*, *C. p. parvum* and *C. p. anthroponosum* (Supplementary Figure S3). However, 33 amino acid residues (highlighted in blue in Figure 3A) are shared by *C. p. anthroponosum* and *C. hominis*, but differ from those present in *C. p. parvum*. These residues are predominantly surface-exposed, supporting the possibility that the two human-associated lineages use this protein to engage host molecules differently from the zoonotic *C. p. parvum*.

**Figure 3.**
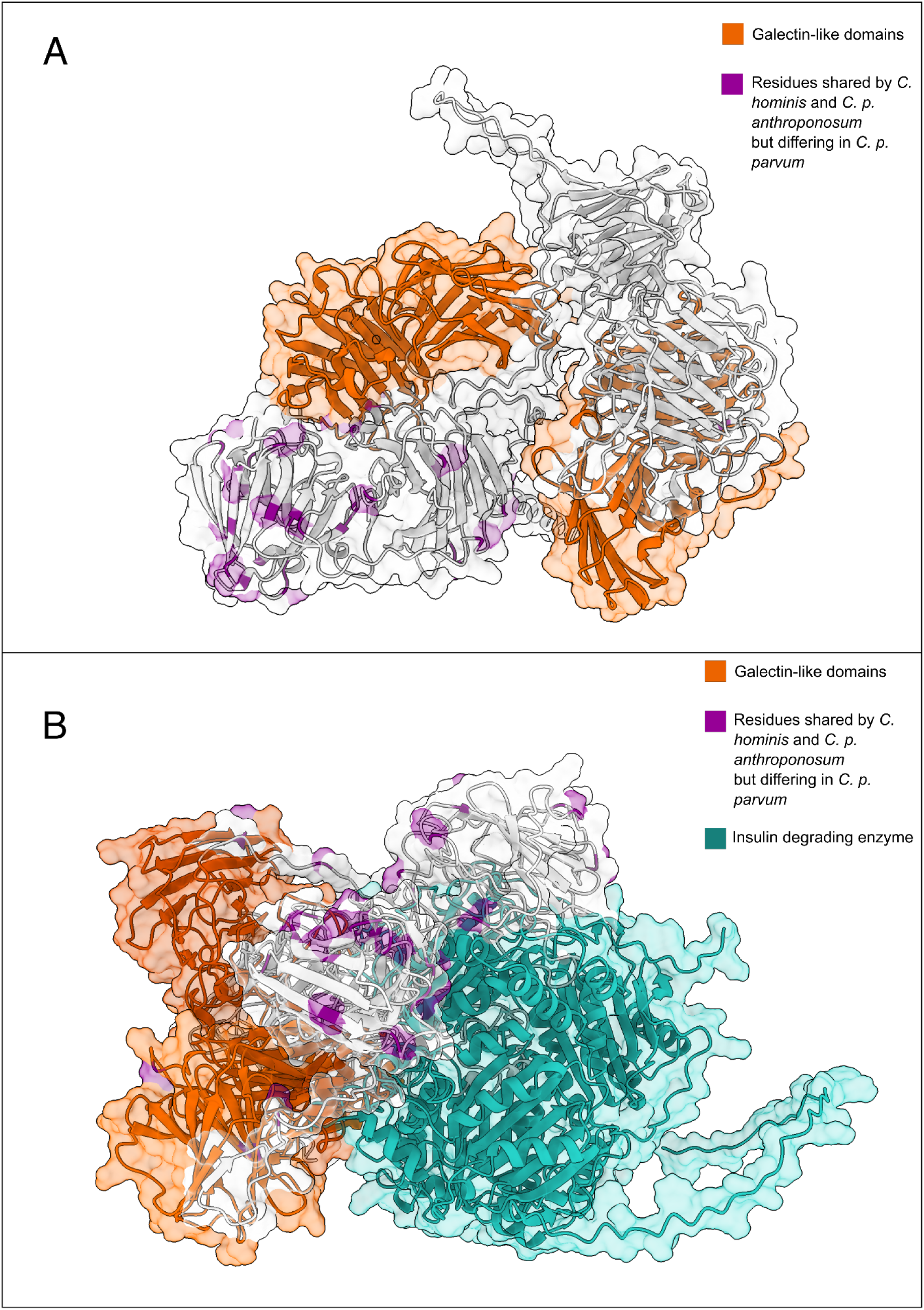
Predicted three-dimensional structure of the *C. p. anthroponosum* galectin-like protein shown as a monomer (A) and in complex with the human insulin-degrading enzyme (B). The backbone of the galectin-like protein is depicted as a light gray cartoon with an overlaid semi-transparent surface. Galectin-like domains are highlighted in dark orange. Residues shared by *C. hominis* and *C. p. anthroponosum* but differing in *C. p. parvum* are shown in purple. In (B), the insulin-degrading enzyme is shown in teal, depicted as a cartoon with an overlaid semi-transparent surface.

### The *Cryptosporidium parvum anthroponosum* galectin-like protein forms a complex with the human insulin-degrading enzyme

On this basis, to investigate the possible role of the introgressed galectin-like protein in the human tropism of *C. p. anthroponosum*, we tested *in silico* whether it could directly interact with human proteins expressed in the intestine, the primary site of parasite colonization ^4^. Out of the 15,187 components of the human intestinal proteome, AlphaFold-Multimer predictions (Supplementary Figure S4) allowed us to identify a single confidently supported interactor of the *C. p. anthroponosum* galectin-like protein (pDockQ = 0.723) (Figure 3B).

This predicted human interactor (accession: P14735) is annotated as “insulin-degrading enzyme” (IDE), namely a zinc metalloprotease that regulates the processing and degradation of insulin and other metabolic peptides ^47^. This interaction is compatible with the observation that, in murine models, galectins can positively influence insulin levels by modulating the activity of proteins involved in its regulation and clearance ^48^. Accordingly, and considering that high human insulin levels were detected in association with increased growth of parasites ^49,50^, *Cryptosporidium* may exploit their galectin-like protein to modulate the host IDE activity and increase their own proliferation. The metabolic context of each host species may further shape the impact of these effects, considering that, unlike humans, ruminants rely less on glucose and insulin signaling pathways ^51,52^. This difference could generate host-specific selective pressures on the galectin-like protein of *Cryptosporidium* spp., which would in turn explain its putative involvement in the host specificity, particularly in the selective adaptation of *C. p. anthroponosum* to humans.

## Conclusions

This work substantiates and expands previous findings on the origin and evolution of *C. p. anthroponosum*^7^, and provides a novel hypothesis for the genetic and functional foundation for its host tropism. We demonstrate that *C. p. anthroponosum* is monophyletic, forming the sister lineage of the zoonotic subspecies *C. p. parvum*. Our analyses further highlighted an introgression between *C. hominis* and *C. p. anthroponosum*, involving a gene encoding for a galectin-like protein. *In silico* predictions revealed a high-confidence interaction of this protein with a key regulator of human insulin metabolism. Since insulin regulation has been implicated in host-parasite interplays ^49,50^, this could represent a credible molecular basis linking the introgression to the human tropism of *C. p. anthroponosum*.

Therefore, our results identify a strong candidate case of introgression linked to host adaptation in *Cryptosporidium*, highlighting the notion that such events have relevant impacts on parasites, shaping host specificity ^53,54^ and, more broadly, host adaptation ^55,56^. Although we did not find evidence of positive selection on the recipient lineage, this would not necessarily be expected if the adaptive event primarily involved the acquisition and retention of an already divergent functional haplotype.

Further exploration of *Cryptosporidium*–host interactions, particularly through experimental validation, will be critical to fully understand the molecular determinants of the host tropism of this parasite and its evolution, and, more in general, of its host interactions, thus paving the way for targeted treatment.

## Supplementary Material

Supplementary Table S1. List of *Cryptosporidium* samples used in this study. The table includes sample ID along with metadata where available such as species/sublineage, subtype, geographic origin, year of isolation, publication and reads accession number.

**Supplementary Figure S1.**
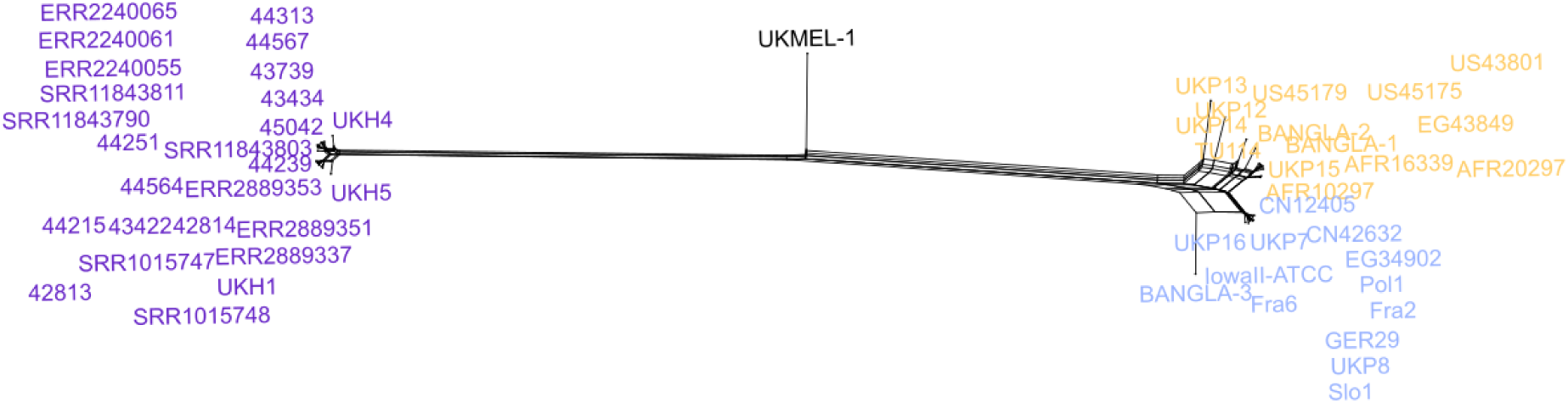
Genome-wide split network showing reticulation between *C. p. anthroponosum* (yellow) and both *C. p. parvum* (blue) and *C. hominis* (purple)

**Supplementary Figure S2.**
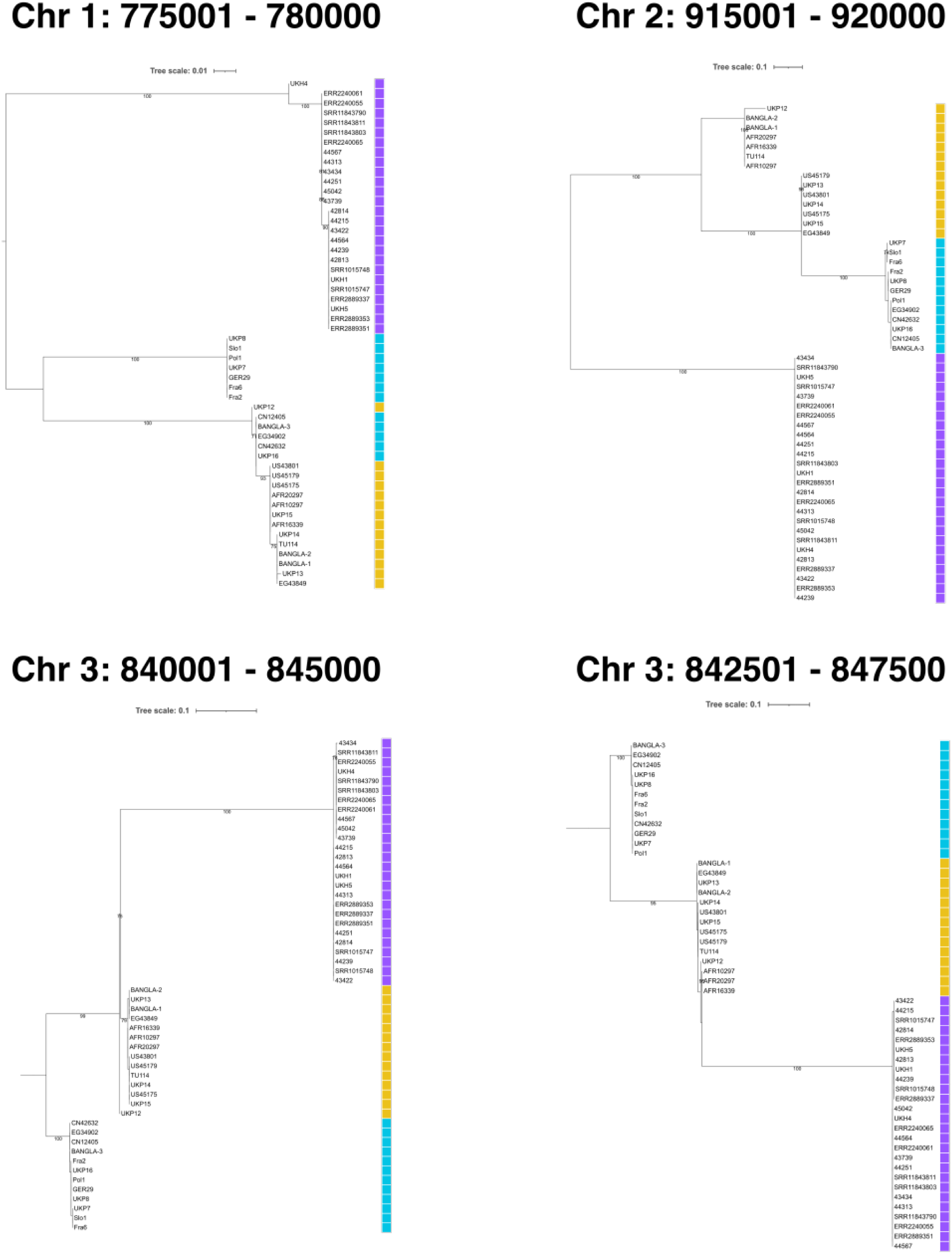
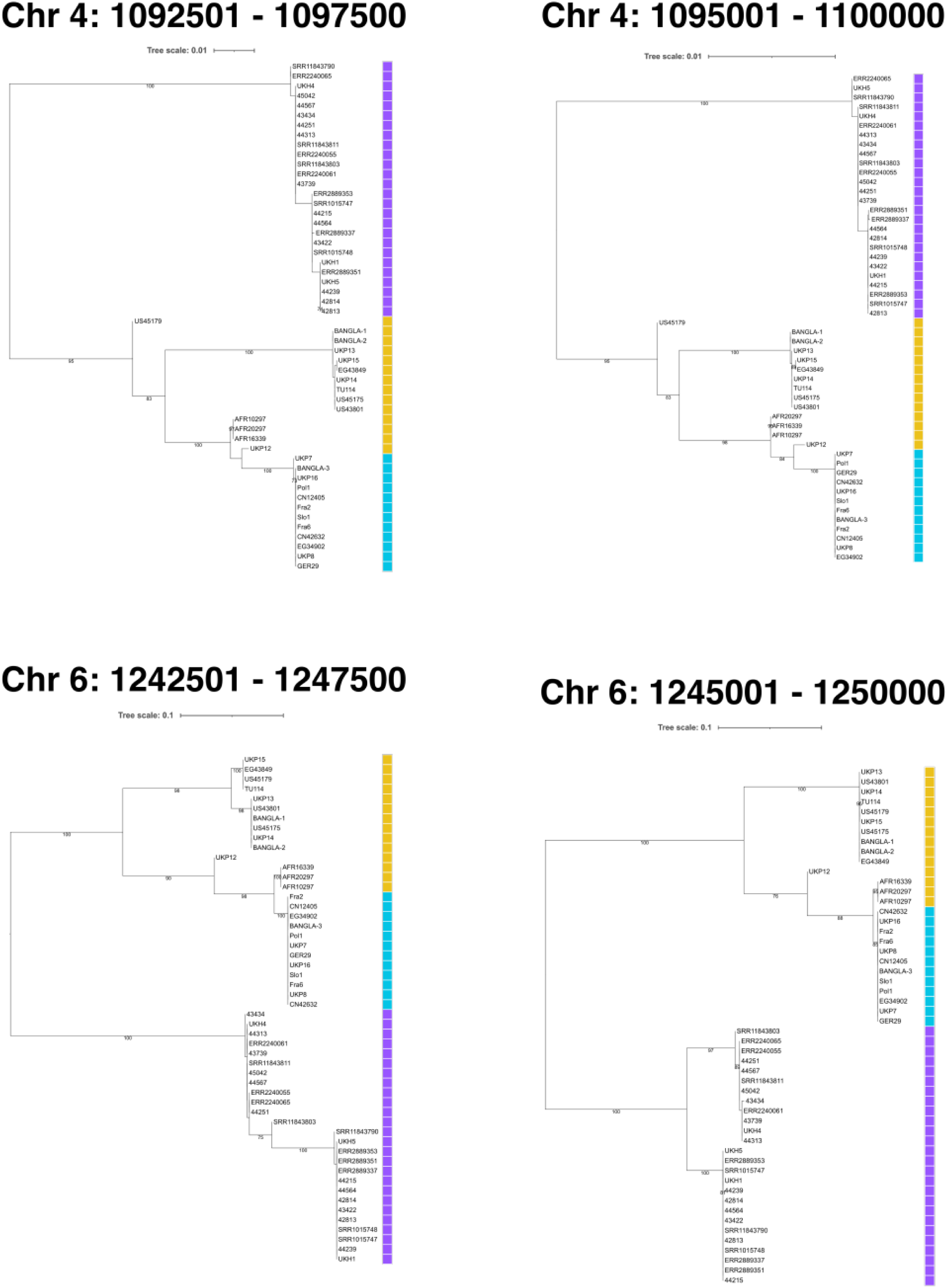
Phylogenies of the candidate introgressed regions, identified through ABBA–BABA analysis as showing elevated fD values in *C. p. anthroponosum*. The colored strip alongside the three correspond to species/sublineages: *C. p. parvum* in blue, *C. p. anthroponosum* in yellow, and *C. hominis* in purple.

**Supplementary Figure S3.**
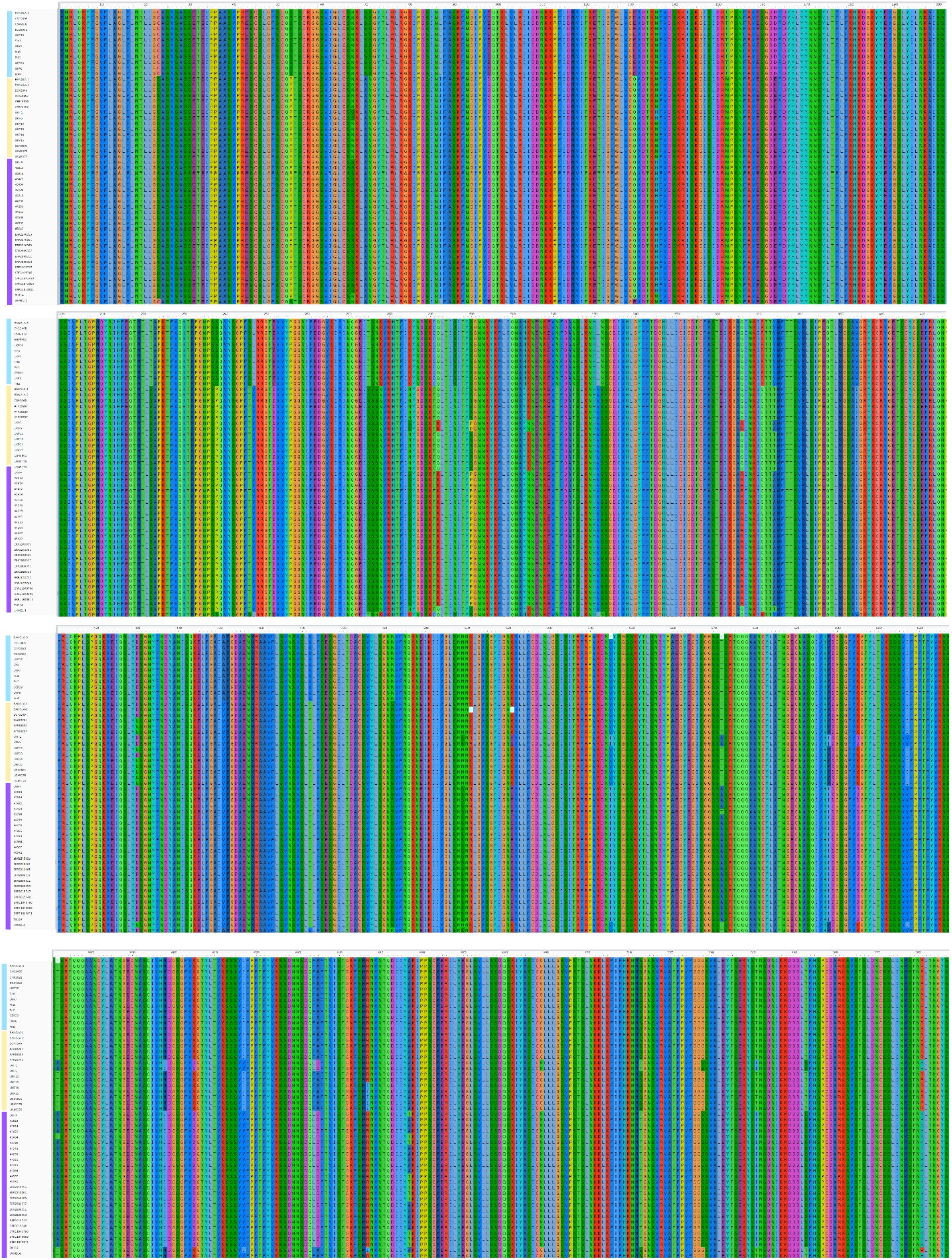
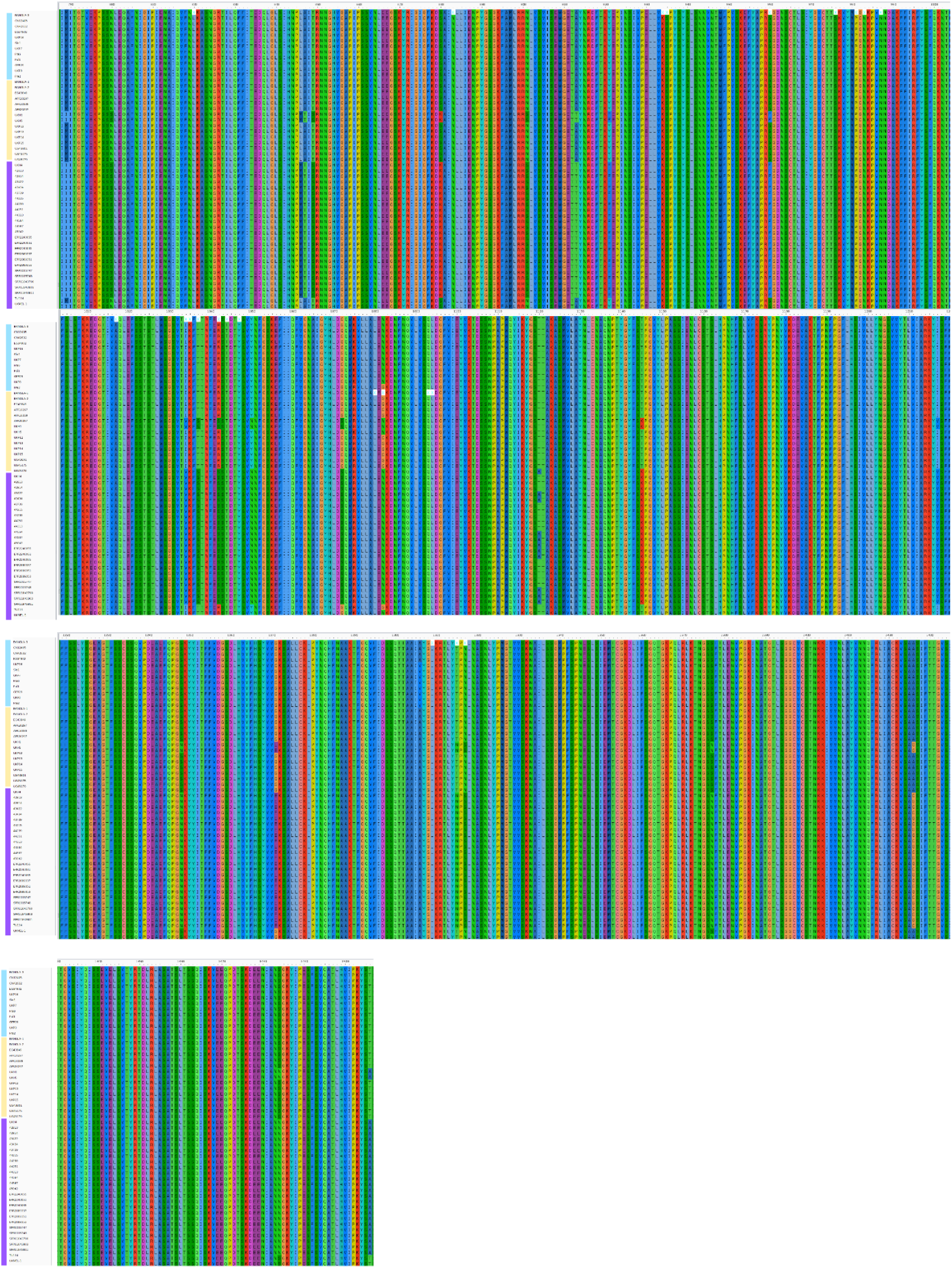
Amino-acid sequence alignment of all sequences obtained from all samples in the dataset for the protein with accession QOY42690.1 from *C. parvum* IOWAII-ATCC reference genome. Each row represents one sequence included in the analysis. A vertical colored strip adjacent to each sequence indicates the species assignment: yellow for *C. p. anthroponosum* (C.p.a), blue for *C. p. parvum* (C.p.p), and purple for *C. hominis.* The alignment allows visualization of sequence conservation and variation across samples and species.

**Supplementary Figure S4.**
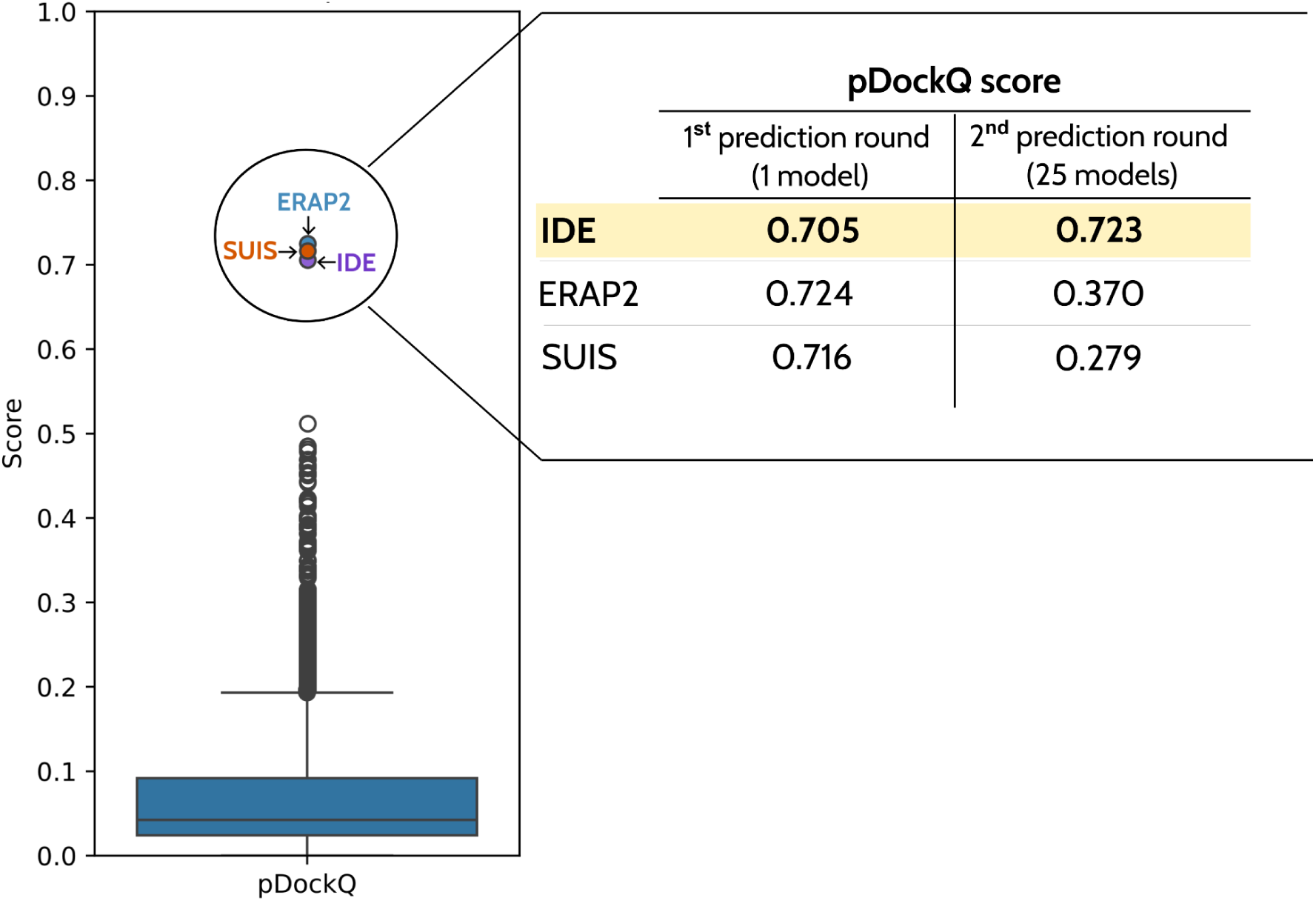
Distribution of pDockQ scores (range 0–1, where 0 indicates an unlikely interaction and 1 a highly likely interaction) obtained from an AlphaFold-Multimer–based screen of predicted interactions between the galectin-like protein of *C. p. anthroponosum* and 15,187 human proteins expressed in the intestine. The initial screening generated a single model for each putative protein–protein complex. The boxplot shown on the left illustrates that the vast majority of predicted complexes cluster at very low pDockQ values (mean = 0.0621), as expected for non-interacting protein pairs. Only three proteins exhibited substantially higher pDockQ scores (∼0.7). These candidates correspond to endoplasmic reticulum aminopeptidase 2 (ERAP2), sucrase-isomaltase (SUIS), and insulin-degrading enzyme (IDE). To reduce the number of potential false positives, a second round of AlphaFold-Multimer predictions was performed, increasing the number of models to up to 25 per protein–protein pair. The table on the right reports pDockQ values for both the initial and refined predictions of these three candidates. Following refinement, pDockQ scores for ERAP2 and SUIS dropped below 0.4, whereas IDE remained the only candidate displaying consistently high pDockQ values, supporting a robust predicted interaction.

